## Supplementary material for "Lymph Node Morphology in Stage II Colorectal Cancer": sup. Table 1

**SUPPLEMENTARY TABLE 1:** Demographics, clinicopathological features and miR-21 expression

| **Variable** |  | **miR-21 < 2 (n=10)** | **miR-21 ≥ 2 (n=9)** | **P-value** |
| --- | --- | --- | --- | --- |
| **Age** | Above Mean (66)  Below | 5  5 | 5  4 | >0.99 |
| **Gender** | Female  Male | 5  5 | 7  2 | 0.35 |
| **T stage** | T3  T4a/b | 6  4 | 7  2 | 0.63 |
| **Histological type** | Adenocarcinoma  Mucinous adenocarcinoma | 9  1 | 7  2 | 0.58 |
| **Grade (differentiation)** | High  Moderate – low | 5  5 | 6  2 | 0.37 |
| **CEA^a^** | <3.5  ≥3.5 | 5  5 | 5  4 | >0.99 |
| **Lymphatic Invasion** | Yes  No | 3  7 | 0  9 | 0.21 |
| **dMMR^b^** | Evidence  No evidence | 6  4 | 2  7 | 0.17 |
| **Multiple polyps** | Yes  No | 4  6 | 5  3 | 0.64 |
| **TDLNs examined for cancer^c^** | ≥12  <12 | 9  1 | 8  1 | >0.99 |

^a^CEA level 3.5 µg/L cut-off (private communication).

^b^Deficiency in at least one of MSH2, MSH6, PMS2, MLH1.

^c^<12 resected lymph nodes is associated with a worse outcome (2).

Tumour miR-21 levels were grouped into miR-21 levels < 2 fold change and miR-21 ≥ 2 fold change. P-values correspond to comparison of miR-21 < 2 and miR-21 ≥ 2 (Fisher’s exact test). Abbreviations: CEA, carcinoembryonic antigen; MMR, mismatch repair; TDLNs, tumour-draining lymph nodes. CEA cut-off value of 3.5ug/L determined through private communication. MMR deficiency based in at least one of MSH2, MSH6, PMS2, MLH1.
