## Supplementary material for "Lymph Node Morphology in Stage II Colorectal Cancer": sup. Table 2

**SUPPLEMENTARY TABLE 2:** Demographics, clinicopathological features and TDLN histomorphology

|  | F density | | GC density | | Primary Follicle density | | F size | | GC size | | Marginal zone | |
| --- | --- | --- | --- | --- | --- | --- | --- | --- | --- | --- | --- | --- |
|  | Mean | p value | Mean | p value | Mean/Median | p value | Median | p value | Median | p value | Mean/Median | p value |
| Female | 13 ± 3 | 0.48 | 9 ± 4 | 0.63 | 5 ± 3 | 0.79 | 0.097 (0.091-0.210) | 0.55 | 0.036 (0.030-0.092) | 0.59 | 0.068 (0.0573-0.101) | 0.75 |
| Male | 12 ± 4 |  | 8 ± 3 |  | 4 ± 3 |  | 0.14 (0.102-0.150) |  | 0.052 (0.033-0.069) |  | 0.086 (0.064-0.089) |  |
| Age >66 | 13 ± 4 | 0.78 | 9 ± 4 | 0.45 | 3 (3-5) | 0.25 | 0.095 (0.084-0.211) | 0.56 | 0.036 (0.033-0.110) | 0.51 | 0.063 (0.050-0.101) | 0.47 |
| ≤66 | 13 ± 3 |  | 8 ± 3 |  | 5 (3-6) |  | 0.12 (0.095-0.251) |  | 0.040 (0.030-0.062) |  | 0.070 (0.064-0.088) |  |
| T3 stage | 14 ± 3 | **0.03** | 9 ± 3 | 0.31 | 5 ± 3 | 0.17 | 0.100 (0.091-0.143) | 0.40 | 0.038 (0.033-0.055) | 0.64 | 0.073 ± 0.018 | 0.17 |
| T4a/b stage | 11 ± 3 |  | 7 ± 4 |  | 4 ± 2 |  | 0.140 (0.095-0.214) |  | 0.072 (0.030-0.111) |  | 0.089 ± 0.035 |  |
| Adenocarcinoma | 13 ± 3 | 0.87 | 9 ± 4 | 0.48 | 4 ± 2 | 0.21 | 0.120 (0.090-0.201) | 0.76 | 0.045 (0.034-0.100) | 0.15 | 0.081 ± 0.028 | 0.35 |
| Mucinous | 13 ± 4 |  | 8 ± 4 |  | 6 ± 4 |  | 0.098 (0.095-0.129) |  | 0.030 (0.027-0.061) |  | 0.068 ± 0.002 |  |
| Well differentiated | 13 ± 4 | 0.51 | 9 ± 4 | 0.81 | 5 ± 3 | 0.58 | 0.098 (0.091-0.140) | 0.35 | 0.037 (0.032-0.072) | 0.62 | 0.065 (0.055-0.090) | 0.06 |
| Moderate- poorly differentiated | 12 ± 3 |  | 9 ± 2 |  | 4 ± 2 |  | 0.150 (0.107-0.210) |  | 0.062 (0.031-0.092) |  | 0.088 (0.077-0.118) |  |
| CEA <3.5 µg/L | 12 ± 3 | 0.15 | 8 ± 3 | **0.04** | 5 ± 2 | 0.35 | 0.100 (0.092-0.140) | 0.51 | 0.037 (0.031-0.052) | 0.38 | 0.067 (0.058-0.089) | 0.65 |
| CEA ≥3.5 µg/L | 14 ± 3 |  | 10 ± 3 |  | 4 ± 3 |  | 0.140 (0.091-0.212) |  | 0.062 (0.032-0.110) |  | 0.068 (0.063-0.100) |  |
| Lymphovascular or peri-neural invasion | 14 ± 4 | 0.18 | 9 ± 3 | 0.82 | 6 ± 3 | 0.13 | 0.098 (0.082-0.133) | 0.24 | 0.036 (0.031-0.045) | 0.40 | 0.068 (0.049-0.090) | 0.35 |
| No invasion | 12 ± 3 |  | 9 ± 4 |  | 4 ± 2 |  | 0.12 (0.093-0.210) |  | 0.047 (0.031-0.100) |  | 0.069 (0.064-0.100) |  |
| dMMR | 14 ± 4 | 0.09 | 10 ± 4 | 0.06 | 4 ± 3 | 0.72 | 0.140 (0.098-0.210) | 0.19 | 0.050 (0.034-0.092) | 0.31 | 0.085 ± 0.025 | 0.31 |
| No dMMR | 12 ± 3 |  | 8 ± 3 |  | 5 ± 2 |  | 0.094 (0.091-0.150) |  | 0.036 (0.031-0.060) |  | 0.073 ± 0.025 |  |
| Multiple polyps | 14 ± 2 | 0.21 | 9 ± 3 | 0.91 | 6 ± 3 | 0.11 | 0.101 (0.093-0.217) | 0.47 | 0.038 (0.027-0.102) | 0.91 | 0.081 ± 0.026 | 0.60 |
| No polyps | 12 ± 4 |  | 9 ± 4 |  | 4 ± 2 |  | 0.110 (0.086-0.140) |  | 0.038 (0.032-0.067) |  | 0.075 ± 0.026 |  |
| Tumour miR-21 ≥2 fold change | 12 ± 3 | 0.80 | 8 ± 4 | 0.43 | 5 ± 2 | 0.24 | 0.095 (0.091-0.102) | **0.02** | 0.030 (0.027-0.039) | **0.01** | 0.065 (0.057-0.069) | **0.01** |
| Tumour miR-21 <2 fold change | 13 ± 3 |  | 9 ± 3 |  | 4 ± 2 |  | 0.150 (0.132-0.230) |  | 0.062 (0.045-0.113) |  | 0.090 (0.086-0.118) |  |

Values expressed as mean ± standard deviation or median ± interquartile range. Unpaired t-tests was used to detect differences between normally distributed continuous data and Mann-Whitney U tests for non-normally distributed data.
