## Supplementary material for "Lymph Node Morphology in Stage II Colorectal Cancer": sup. Figure 1

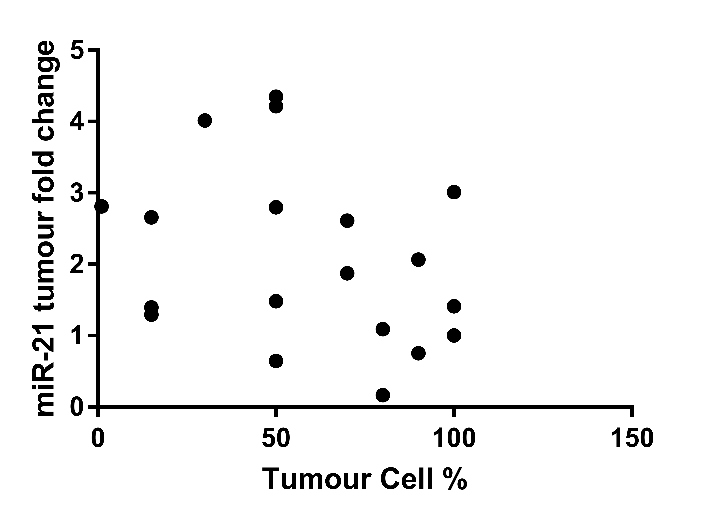

***SUPPLEMENTARY FIGURE 1:*** *Correlation of miR-21 tumour fold change and tumour cell percentage.* *r^2^ = 0.094, p = 0.20, Pearson correlation.*
